# Imaging Membrane Curvature inside a Model Immunological Synapse in RBL-2H3 Cells using TIRF Microscopy with Polarized Excitation

**DOI:** 10.1101/608463

**Authors:** Rosa Machado, Justin Bendesky, Madison Brown, Kathrin Spendier, Guy M. Hagen

**Author notes:** Correspondence (K.S.); (G.H.).

## Abstract

Total internal reflection fluorescence microscopy with polarized excitation (P-TIRF) can be used to image nanoscale curvature phenomena in live cells. We used P-TIRF to visualize rat basophilic leukemia cells (RBL-2H3 cells) coming into contact with a supported lipid bilayer, modeling an immunological synapse. These studies help correlate the dynamics of cell surface molecules with the mechanical properties of the plasma membrane during synapse formation.

## 1. Introduction

The ability of eukaryotic cells to communicate with each other is important for numerous biological processes, including cell growth, motility, and immune function. This communication can occur through soluble factors (cytokines, interleukins, etc.), or by physical cell-cell contact. In the latter case, the immunological synapse is one of the most important communication strategies in immune cells. These intimate cell-cell contacts result in intracellular signaling and is accompanied by large-scale spatial reorganization of membrane proteins within the cell-cell junction. For example, the ability to form a synapse is a well-known communication strategy between T cells and antigen-presenting cells such as B-cells[1,2]. A recent study by Carroll-Portillo and collaborators have shown that mast cells can form a synapse with dendritic cells, and that this can facilitate antigen transfer in T cell activation processes[3]. This suggests to us that mast cells and basophils may play a larger roles in signaling between physically contacting cells.

To study immunological synapse formation we used total internal reflection fluorescence (TIRF) microscopy of fluorescently labeled cells interfacing with a supported lipid bilayer (SLB) containing an appropriate ligand[4]. We have previously demonstrated that rat basophilic leukemia (RBL-2H3) cells possess the machinery to form an immunological synapse in response to antigen-containing SLBs[4–6]. RBL-2H3 cells are derived from rat basophils, but are commonly used in studies of allergic responses which are typically associated with mast cells[7,8]. Similar to mast cells, widely used RBL-2H3 cells degranulate in response to multivalent antigens which crosslink immunoglobulin E (IgE) when bound to its high affinity receptor, FcεRI[8,9]. RBL-2H3 cells can be weakly activated when in contact with fluid lipid bilayers containing monovalent ligands such as dinitrophenol (DNP)[4,10]. In this model system IgE-FcεRI are not cross-linked but aggregate at cell surface protrusions that form the initial contact points with the substrate[6]. After initial IgE-FcεRI cluster formation, small receptor clusters coalesce to form a large central patch, termed the mast cell synapse[4–6]. During activation, RBL-2H3 cells are known to change their morphology, with the cell becoming more ruffled[11]. However a detailed picture of how the transformation of cell membrane shape affects the organization of receptors is still lacking.

Recent studies, for example those summarized in Ref. [12], have focused on identifying proteins that are involved in membrane curvature generation and sensing. In these investigations a surprising abundance of proteins that couple membrane shape to cellular membrane function have been discovered. In this context, we have previously shown that an elevated concentration of curvature-inducing peptides can interfere with the formation of a mast cell synapse in RBL-2H3 cells[13], and it is known that receptor activity at the plasma membrane can be enhanced by membrane curvature changes[14]. To investigate the role of the plasma membrane environment in the formation of immunological synapses in RBL-2H3 cells, we wish to relate the spatial and temporal dynamics of the IgE receptor, FcεRI, with what may be very small changes in cell membrane topography. This can be accomplished using polarized-excitation TIRF microscopy (P-TIRF)[15–24].

We used P-TIRF to determine the relative molecular orientation of the fluorescent lipophilic tracer DiI-C_16_ in RBL-2H3 cells which were labeled with fluorescent IgE. DiI-C_16_ inserts into the plasma membrane of cells in a specific orientation such that the chromophore is always oriented parallel to the membrane. The dye is preferentially excited by linearly polarized light according to its orientation. DiI-C_16_ molecules in areas of the membrane with high curvature in the axial direction are preferentially excited by P-polarized excitation[21]. In a microscope, P-polarized light can be achieved using through-the-objective type TIRF illumination with high NA objective lenses[21], and a ratio of P-polarized excitation and S-polarized excitation images can be used to assess dye orientation and thereby axial membrane curvature[19].

In this study we used P-TIRF to image live RBL-2H3 cells in which we labeled the membrane receptor FcεRI with fluorescent IgE (IgE-488) and labeled the plasma membrane with DiI-C_16_. We then allowed the cells to settle on and engage supported lipid bilayers containing lipids bearing the IgE ligand DNP. This allows us to correlate receptor patch formation and dynamics with measurements of membrane curvature, while also exploiting the optical sectioning property of TIRF microscopy. We used an innovative optical design and sensitive sCMOS and EMCCD detectors to capture high signal to noise ratio images, which we used to generate P-TIRF ratio images that indicate membrane curvature. These are the first experiments which combine imaging of membrane curvature phenomena with simultaneous imaging of the formation of a model immunological synapse in RBL-2H3 cells that we are aware of.

## 2. Materials and Methods

### 2.1 Samples

RBL-2H3 cells, obtained from ATCC (Manassas, VA), were maintained in DMEM supplemented with 10% FCS, 100 U/ml penicillin, and 100 U/ml streptomycin (all from Invitrogen) at 37 °C, 5% CO_2_, and 100% humidity. Cells were grown in tissue culture flasks, and were transferred to suspension culture petri dishes 24-48 hours before imaging.

We labeled anti-dinitrophenol IgE (anti-DNP-IgE, clone SPE-7, Sigma-Aldrich, D8406) with Dylight 488 NHS ester (Thermo Fisher Scientific, 46403) according to the manufacturer’s protocols (IgE-488). After size-exclusion chromatography, the final dye to protein ratio was typically 5-6 dye molecules per protein molecule as measured with a Nanodrop spectrophotometer (Thermo Fisher Scientific). Cells in suspension culture dishes were labeled with 0.5 nM fluorescent IgE for typically 12-20 hours before imaging.

To label the plasma membrane for P-TIRF measurements, we labeled the cells with 1 μM DiI-C_16_ (Thermo Fisher Scientific, D384). To prepare the DiI-C16 we dissolved the solid material in ethanol at a concentration of 100 mM, then diluted it to 1 μM in PBS. After 5-10 minutes incubation at 37 °C, we washed the cells with fresh PBS.

We formed liposomes containing 75% 1-palmitoyl-2-oleoyl-sn-glycero-3-phosphocholine (POPC) and 25% 1,2-dihexadecanoyl-sn-glycero-3-phosphoethanolamine-N-[6-[(2,4-dinitrophenyl)amino]caproyl] (DNP-CAP-PE, Avanti Polar Lipids) by mixing the two components dissolved in chloroform, then evaporating the solvent with a blow-down of compressed air. We then dissolved the mixture in PBS, and sonicated it on ice for 10 minutes using a probe type sonicator (Branson Sonifier 450), until the liposome mixture appeared clear.

To prepare supported lipid bilayers, we pipetted 50 μl of liposome mixture on to a clean petri dish at 37 °C, then overlaid the droplet with a 25 mm #1.5 coverslip that had been cleaned in piranha solution (30% H_2_O_2_ in concentrated H_2_SO_4_)[25]. After 15 minutes, we immersed the petri dish and coverslip in distilled water, then inverted the coverslip and placed it in a microscope imaging chamber (RC-40LP, Warner Instruments). This prevented the bilayer from coming into contact with air.

### 2.2 Microscope setup and acquisition

Our setup is based on a DMI300B or DMI8 microscope with an oil immersion HCX PL APO 100×/1.47 NA TIRF objective (Leica, Manheim, Germany). We used an Evolve 512 EMCCD camera (Photometrics, Tucson, Arizona) and micromanager software[26] or a Zyla 4.2+ sCMOS camera with IQ software (Andor, Belfast, Northern Ireland). The TIRF module (custom designed by Spectral Applied Research, Ontario, Canada) uses a liquid crystal polarization rotator and controller (Meadowlark Optics, Frederick, Colorado). The controller is signaled to switch polarizations using an Arduino under the control of micromanager software[27]. The time required to switch polarization states is less than 10 ms, allowing us to record pairs of images with P-polarized and S-polarized excitation with very little delay between the images, and with no moving parts.

Sample fluorescence was isolated with a DV2 image splitter (Photometrics) or with a Lambda 10-B filter wheel (Sutter Instrument, Novato, California) and emission filters appropriate for IgE-488 and DiI-C_16_ (ET series, Chroma, Bellows Falls, Vermont). For live cell experiments with SLBs, we warmed the objective to 37 °C using an objective heater (Bioptechs). In some experiments we used a custom-made microscope incubation chamber. A diagram and photograph of the microscope setup are shown in Figure 1.

This setup uses a small right-angle prism (legs ~2mm) to steer the laser beam into the back aperture of the objective[28]. A second small right-angle prism steers the reflected beam into a beam dump. This approach eliminates the need for a dichroic mirror. The two prisms, and also one lens, are mounted on stepper-motor actuated stages under computer control. This allows precise and reproducible positioning of the laser beam in the objective back aperture, thereby allowing us to achieve the desired incidence angle (TIRF angle), and therefore the desired penetration depth of the evanescent field. A rotating plate is also stepper motor actuated under computer control and is used for wavelength compensation such that different illumination wavelengths can have the same evanescent field penetration depth. In this setup the azimuthal position of the beam in the back aperture is fixed. This can result in uneven illumination and interference fringes in the image, but this problem was not severe in our case. Spinning the laser beam around the periphery of the back aperture within a single camera exposure can help eliminate such interference patterns[29].

**Figure 1.**
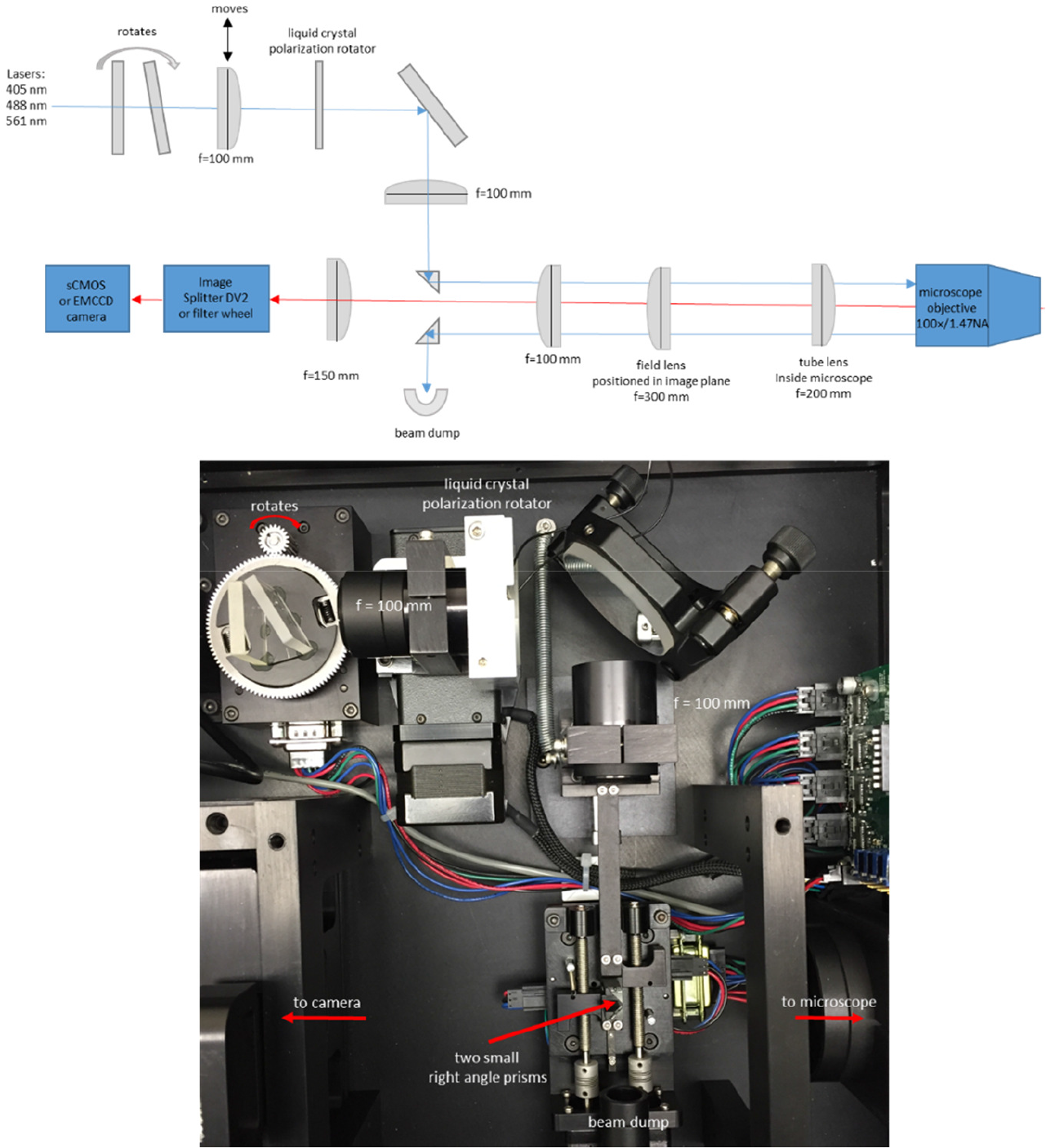
P-TIRF microscope setup. Diagram and photograph. Diagram not to scale. Use of small (~2 mm) right-angle prisms[28] to steer the laser beam removes the need for dichroic mirrors. The moving lens and prisms are stepper motor actuated under computer control and allow precise adjustment of the beam position, and thus of the penetration depth of the evanescent field.

### 2.3 Membrane Curvature Visualization

A detailed theoretical discussion of the workings of P-TIRF can be found in reference [16]. Briefly, a polarization component of light along the optical axis or the z-axis of the microscope is unique to TIRF microscopy. With epi-illumination, the electromagnetic field propagates along the optical axis and only has components in the x-y plane with respect to the sample. With TIRF illumination, P-polarization results in polarized illumination along the x-axis and the z-axis, with the x-component being about 5% of the z-component. S-polarization results in polarized illumination only along the y-axis. The probability of a fluorophore to absorb a photon is calculated by the dot product of the fluorophore’s excitation dipole and the polarization of the absorbed light. Therefore for fluorophores in which the chromophore is oriented in a specific way, one can determine the orientation of the fluorophore by switching between P-polarized and S-polarized TIRF excitation.

The emission light intensities P and S gathered by an objective from P-polarized and S-polarized excitation are[21]

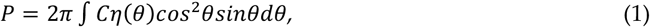

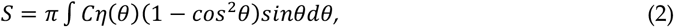

where C = P + 2S is the local concentration of fluorophores, and η(θ) is a factor that describes the orientational distribution of the detected fluorophores. DiI-C_16_ molecules insert in the membrane with their excitation dipoles parallel to the plane of the membrane[15]. DiI-C_16_ molecules that are located in areas of the membrane with high axial curvature are excited preferentially by P-polarized excitation in TIRF and a simple ratio R (or ratio image)

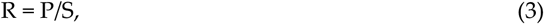

can be calculated to indicate the local orientation of the cell membrane.

To determine if the excitation intensities are the same for both polarizations, we collected signals (using P- or S-polarized epi-illumination) from DiI-C_16_ in solution for both polarizations. We found that this ratio for polarized excitation was close to 1.0. Therefore to calculate the ratio we simply divided the P and S images without any additional processing. In some cases we averaged multiple ratio images to reduce noise.

## 3 Results

### 3.1 Membrane curvature can be detected using P-TIRF and the dye DiI-C_16_

We grew RBL-2H3 cells on coverslips overnight, then labeled them with 1 μM DiI-C_16_. After washing the cells three times with PBS, we imaged them using P-TIRF in PBS or in media in an open-top chamber. We typically recorded 1-40 images in each orientation of polarized excitation, switching the polarization between each acquired image. An example of the results are shown in Figure 2. In this example, large differences are apparent between the two excitation polarizations. Figures 2a and 2b are shown with the same brightness and contrast settings, while Fig. 2c shows a two-color overlay to emphasize the differences. Figure 2d shows an average of 10 ratio images, with the brightness adjusted to a scale ranging from ratios of 0-3. We measured the P/S ratio, Equation (3), in two regions of interest (indicated in Fig. 2d). We found that the ratio was 1.822 ± 0.295 (mean ± standard deviation) in region 1 (high curvature area), and 0.694 ± 0.0661 in region 2 (low curvature, flat area). When creating two-color overlay or ratio images, we measured the lateral shift between the two images using the ‘find shift’ routine within DIPImage[30], an image processing toolbox for MATLAB (The Mathworks, Natick, MA). We found that the shift between the two images was less than about 0.3 pixels in both X and Y. In our case this corresponds to less than 15 nm and so we did not perform an image registration procedure.

**Figure 2.**
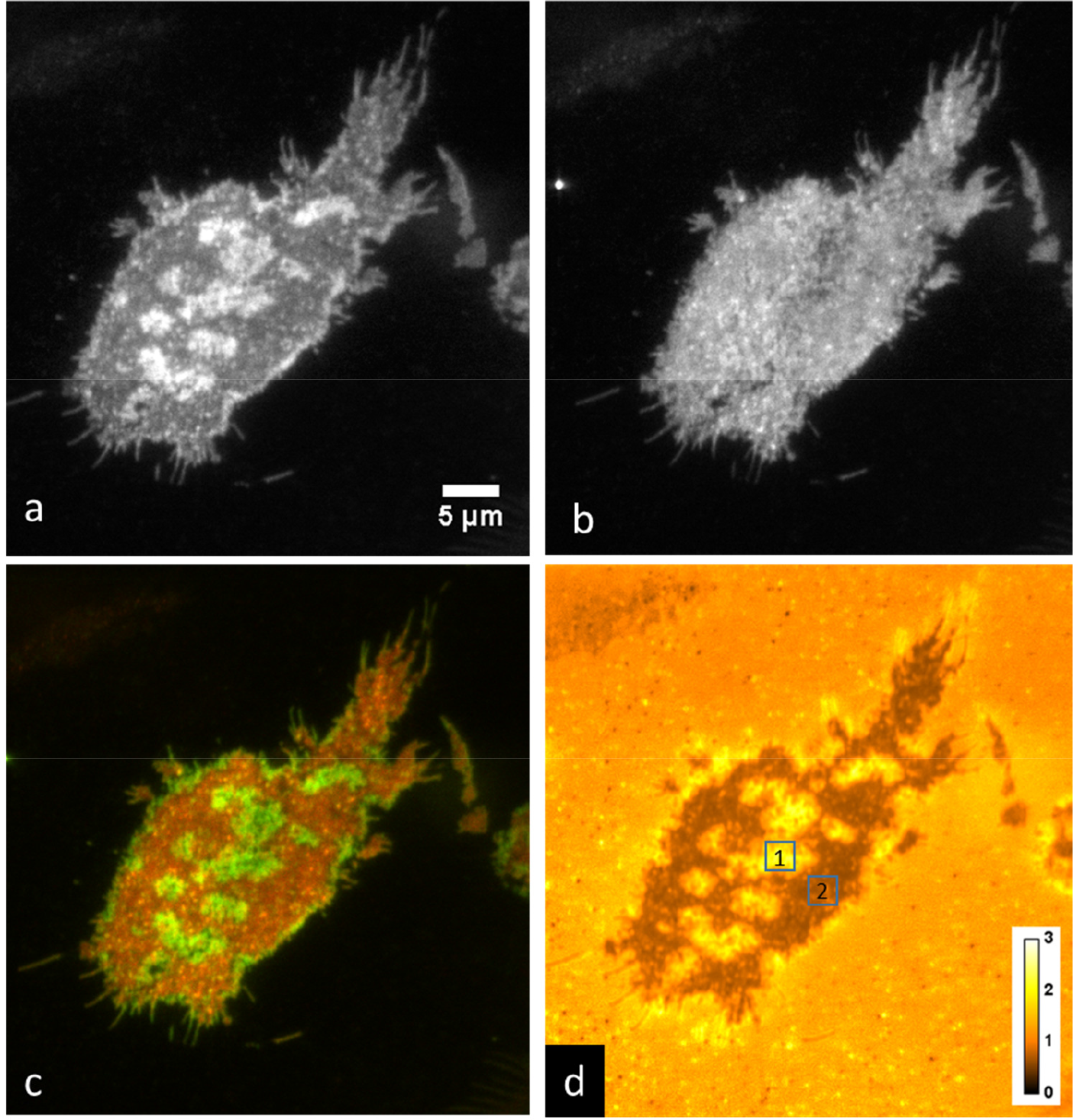
Imaging membrane curvature with DiI-C_16_ in RBL-2H3 cells grown on a coverslip acquired with an Evolve 512 EMCCD camera. Shown are a) P-polarized excitation, b) S-polarized excitation, c) red-green two-color overlay of P-polarized excitation and S-polarized excitation, d) average of ten P/S ratio images. ROI 1: P/S = 1.822 ± 0.295; ROI 2: P/S = 0.694 ± 0.0661.

Axial membrane curvature is high in filopodia, which are approximately cylindrical and thus have a high aspect ratio[24]. We imaged membrane curvature using DiI-C_16_ labeled RBL-2H3 cells grown on a coverslip, acquired with the Zyla 4.2+ sCMOS camera. Figure 3a shows an average of 40 P/S ratio images, while figure 3b shows a zoomed-in view of the region of interest indicated in figure 3a. We measured the P/S ratio from top to bottom along the line indicated in Figure 3b, and plotted the results in the inset in the figure.

**Figure 3.**
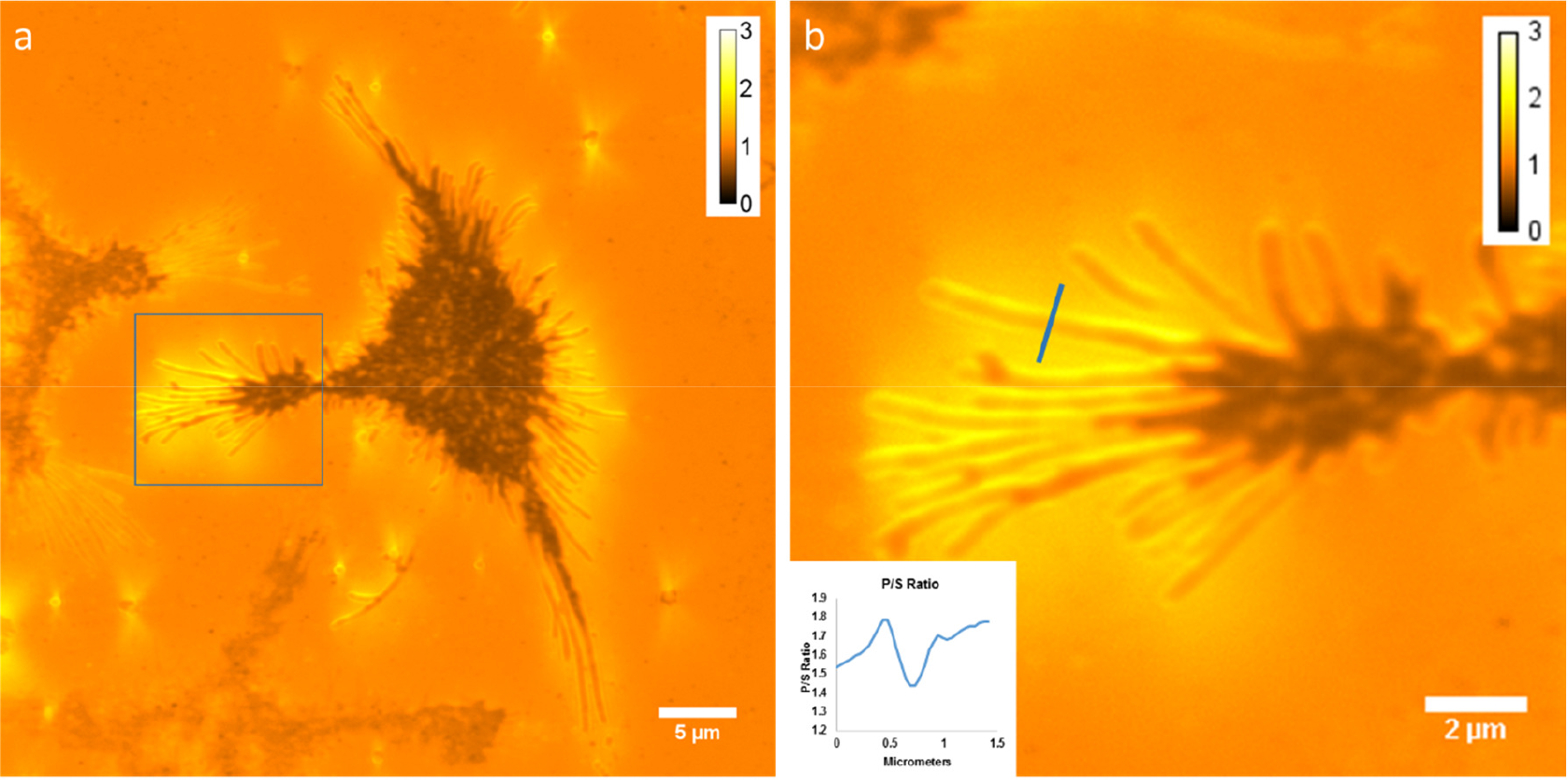
Imaging membrane curvature with DiI-C_16_ in RBL-2H3 cells grown on a coverslip acquired with the Zyla 4.2+ sCMOS camera. a) Average of 40 P/S ratio images. b) zoom of the region indicated in a. The inset shows a plot of the P/S ratio along the indicated line (from top to bottom of the line).

### 3.2 Formation of an immunological synapse on a supported lipid bilayer

We next imaged RBL-2H3 cells that had been labeled overnight with DyLight488-IgE as they came into contact with a supported lipid bilayer containing 25% DNP-CAP-PE. As described above, the IgE-488 is bound to cell surface cell surface IgE receptors (FcεRI), and also to DNP-CAP-PE lipids in the bilayer. After 10-15 minutes, we observed a wide variety of model immunological synapses in the form of dynamic patches of labeled IgE receptor. This is shown in figure 4, a gallery of images with various sizes and shapes of IgE receptor patches.

We tested the fluidity of the supported lipid bilayer using a fluorescence recovery after photobleaching (FRAP) method. To do this we labeled the bilayer with a 1% solution of a BODIPY-conjugated lipid (Thermo Fisher Scientific, D3803), and performed a FRAP experiment using a SP5 confocal microscope (Leica) with warmed objective to 37 °C using an objective heater (Bioptechs). Fitting of the fluorescence recovery to a diffusion model yielded a diffusion coefficient of 1.6 μm^2^/sec. Widefield observations of the BODIPY-labeled bilayer showed that the bilayer was uniform and intact throughout the coverslip surface.

**Figure 4.**
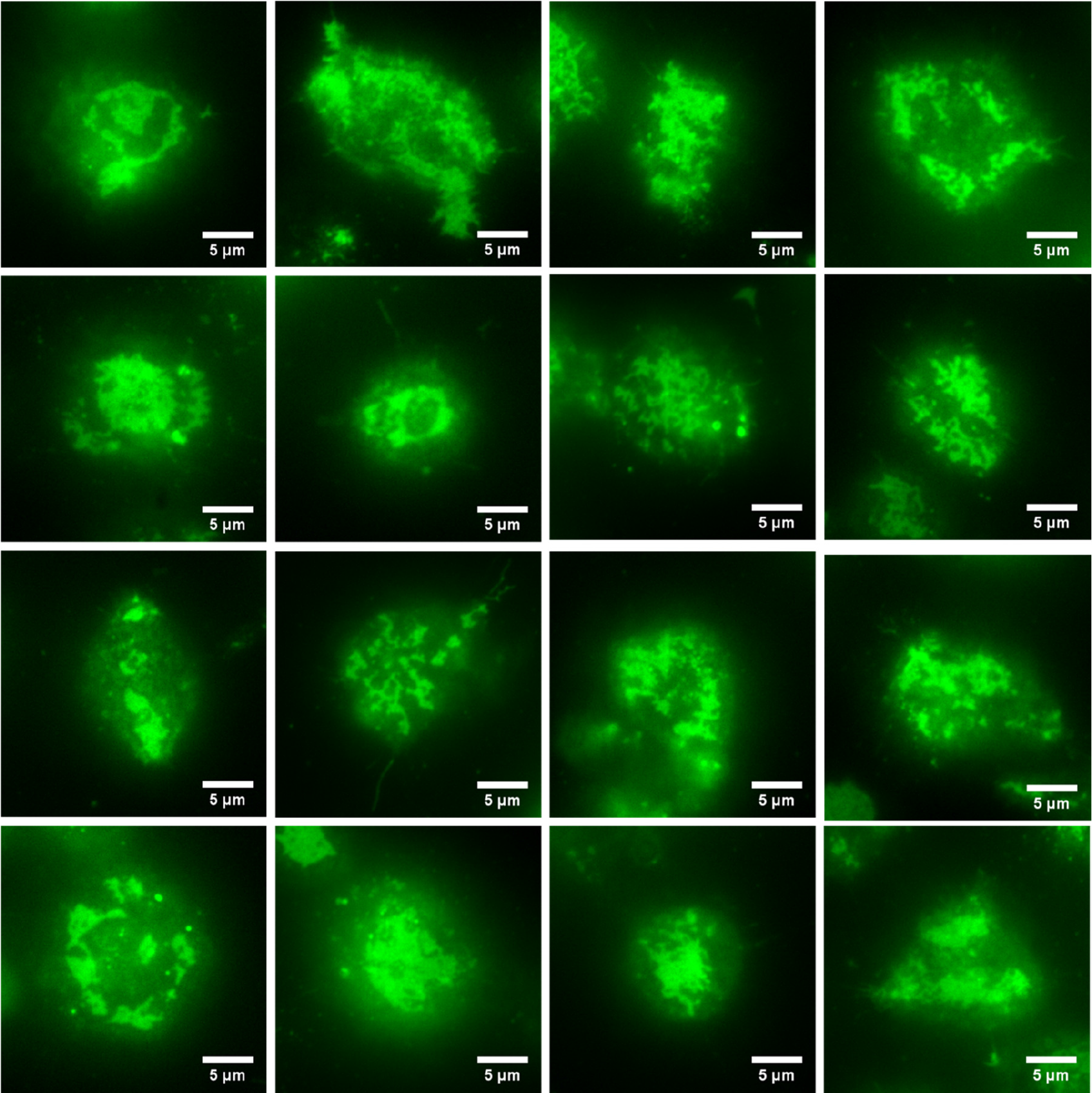
Gallery of images of RBL-2H3 cells labeled overnight with IgE-488 in contact with a supported lipid bilayer containing a lipid derivatized with the IgE ligand DNP. After 10-15 minutes of contact at 37 °C, we observed a wide variety of patches of IgE-488 labeled FcεRI. Images were acquired with the Zyla 4.2+ sCMOS camera.

### 3.3 Imaging of IgE-488 labeled FcεRI with simultaneous imaging of membrane curvature

We next imaged RBL-2H3 cells that had been labeled overnight with IgE-488 and were in contact with a supported lipid bilayer containing a lipid derivatized with the IgE ligand DNP, but in this case the cell membrane was also labeled with DiI-C_16_. This experiment is shown in Figure 5. We observed in many cases that ‘holes’ in the FcεRI patch are evident (for example in Fig 5i), but that these holes are not devoid of cell membrane (Fig 5j). We further analyzed the images by segmenting the image so as to create masks (see supplemental Fig. S1) corresponding to regions containing IgE-bound FcεRI (+IgE) or lacking IgE-bound FcεRI (−IgE), then plotted normalized histograms of the number of pixels with particular P/S ratios in +IgE and −IgE regions (Fig 5d,h,l). Membrane regions containing IgE-bound FcεRI consistently had lower P/S ratios than those regions lacking the receptor, though this varied considerably from cell to cell. For each cell shown in Figure 5, a two-sample Kolmogorov-Smirnov test was performed using the statistics toolbox within MATLAB. For all cells, the test rejected the null hypothesis that P/S ratios in +IgE and −IgE regions come from populations with the same distribution at the 5% significance level.

**Figure 5.**
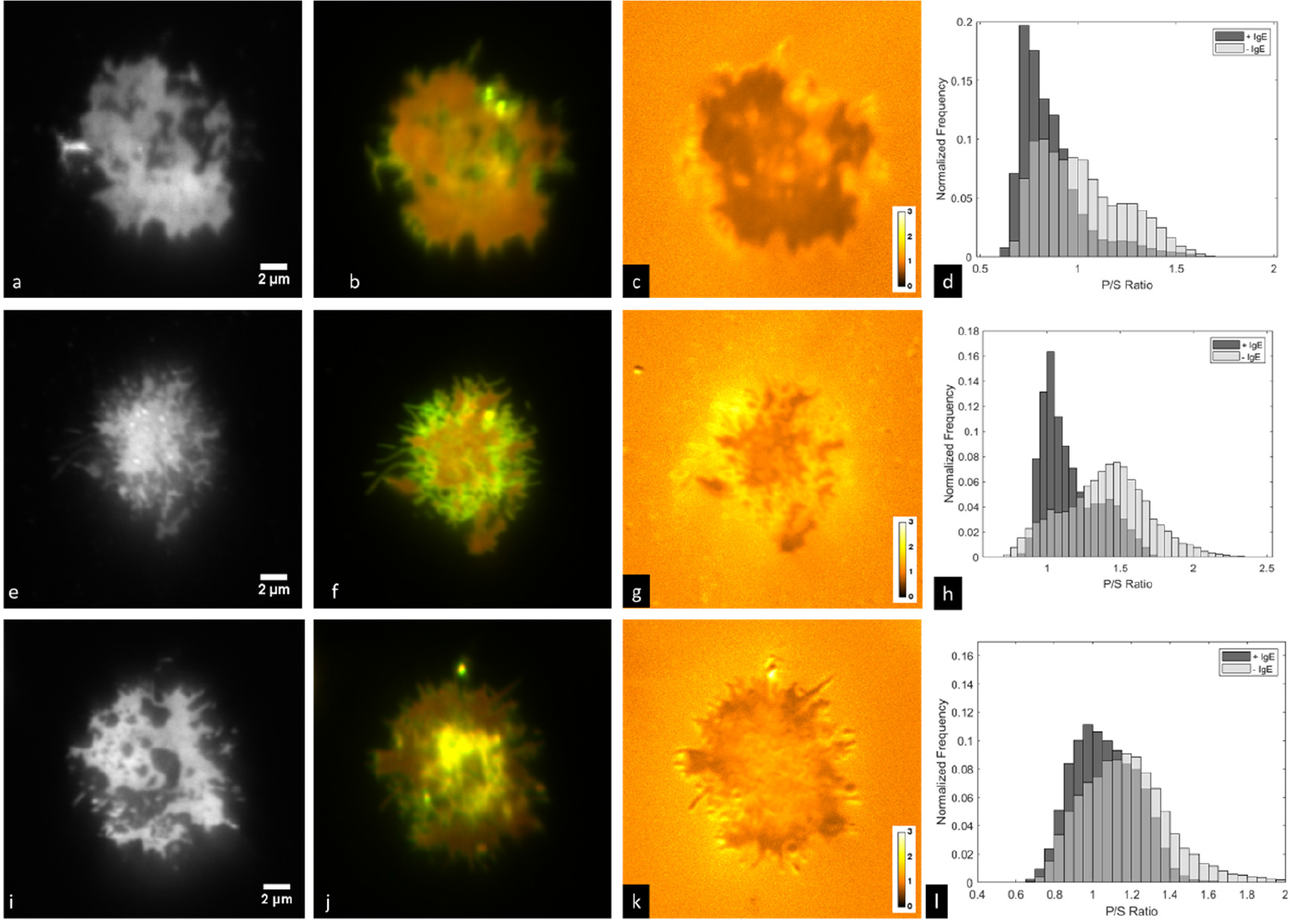
RBL-2H3 cells labeled with IgE-488 and DiI-C_16_ in contact with a supported lipid bilayer containing 25% DNP-CAP-PE. Shown are a, e, i) IgE-488; b, f, j) two-color overlay of P-polarized excitation and S-polarized excitation signals from DiI-C_16_; c, g, k) P/S ratio images of DiI-C_16_ signals; d, h, l)histograms of membrane regions with particular P/S ratios for regions positive (+IgE) or negative (−IgE) for IgE-bound FcεRI. Images were acquired with the Evolve 512 EMCCD camera and DV2 image splitter.

### 3.4 Time-Lapse imaging of a model immunological synapse

Finally, we imaged RBL-2H3 cells in contact with a supported lipid bilayer over the course of about 4 minutes. We observed that the IgE-488 bound FcεRI formed a dynamic patch which continually changed shape. Figure 6 shows time-lapse imaging of IgE-488 bound FcεRI and a two-color overlay of P-polarized excitation and S-polarized excitation signals from DiI-C_16_. 10 time points out of 108 total are shown. Supplementary Video S1 shows the entire image sequence.

**Figure 6.**
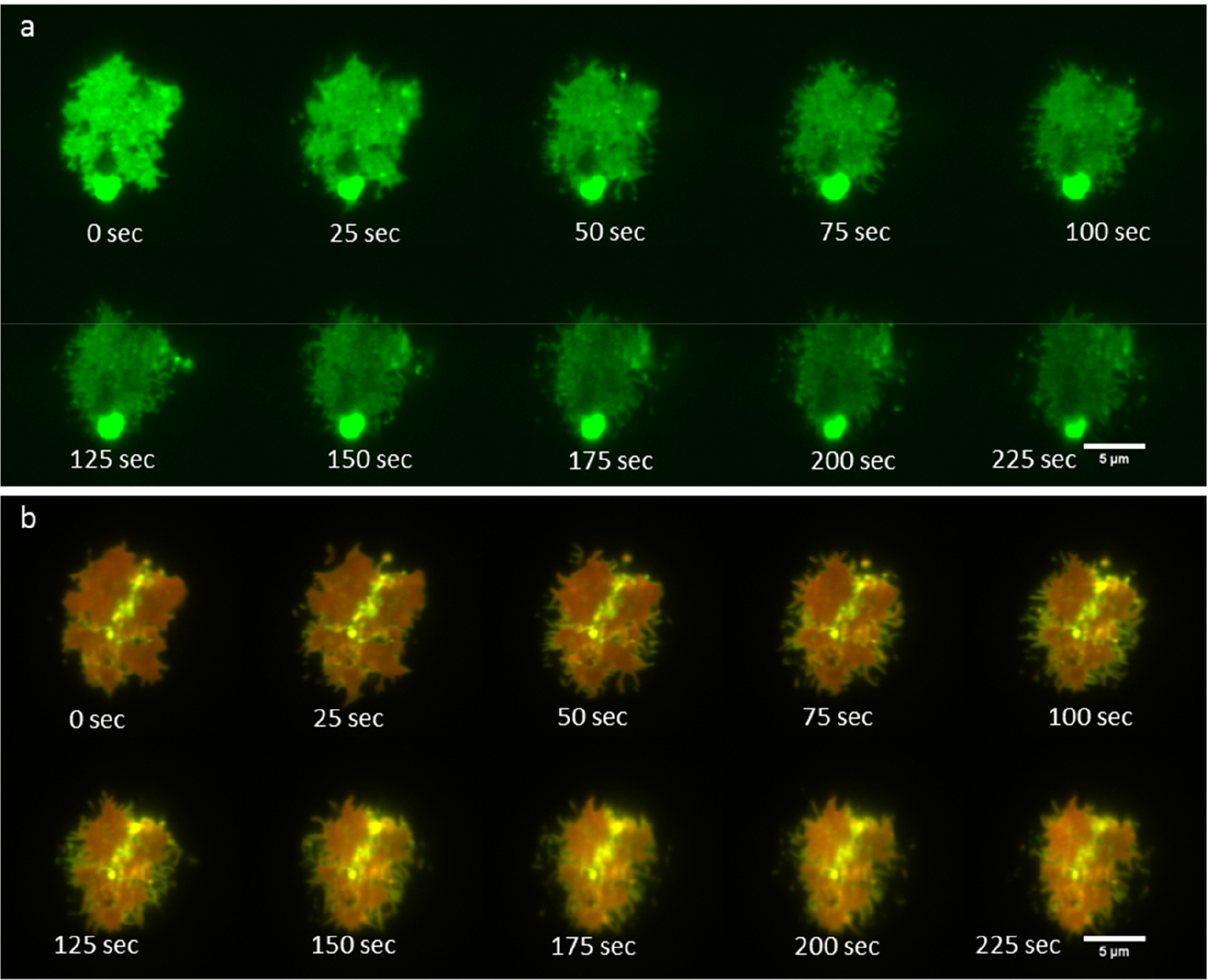
Time-lapse imaging of IgE receptor patch dynamics together with P-TIRF imaging of membrane curvature. a) IgE-488 bound FcεRI; b) P-S two-color overlay image. Images were acquired with the Zyla 4.2+ sCMOS camera.

## 4. Discussion

The relationship between membrane curvature phenomena and the organization of cell surface receptors responsible for signaling in RBL-2H3 cells has not been well explored. Here we used P-TIRF microscopy to image a model of immunological synapse formation in live RBL-2H3 cells. We detected membrane curvature using a fluorescent probe that exhibits a specific orientation in the plasma membrane, and correlated this with the organization and dynamics of the IgE receptor. Membrane regions containing IgE-bound FcεRI consistently had lower P/S ratios than those regions lacking the receptor, indicating that regions lacking the receptor are more curved than regions containing the receptor within the cell-substrate contact zone.

In the future, to help reveal the nature of the curvature features we detect, live cell super-resolution microscopy will be valuable. Structured illumination microscopy[31,32] would allow imaging of receptor patch formation with super-resolution, and single molecule localization microscopy methods[33,34] would allow evaluation of nanoscopic membrane features in relation to IgE receptor dynamics.

## Supplementary Materials

Figure S1: Image masks used to create normalized histograms of the number of pixels with particular P/S ratios in +IgE and −IgE regions shown in Fig. 5. Figure S2. Image showing composite of IgE signal (red) and P-S ratio (green) from a) Figure 5i and 5k and b) Figure 5a and 5c of the main text.

## Author Contributions

Conceptualization, K.S.; Formal analysis, R.M., K.S. and G.M.H; Funding acquisition, K.S. and G.M.H.; Investigation, R.M., J.B., M.B. and G.M.H.; Methodology, K.S. and G.M.H.; Supervision, K.S. and G.M.H.; Visualization, R.M. and G.M.H.; Writing - original draft, K.S. and G.M.H.; Writing - review & editing, K.S. and G.M.H.;

## Supplementary Materials

**Figure S1.**
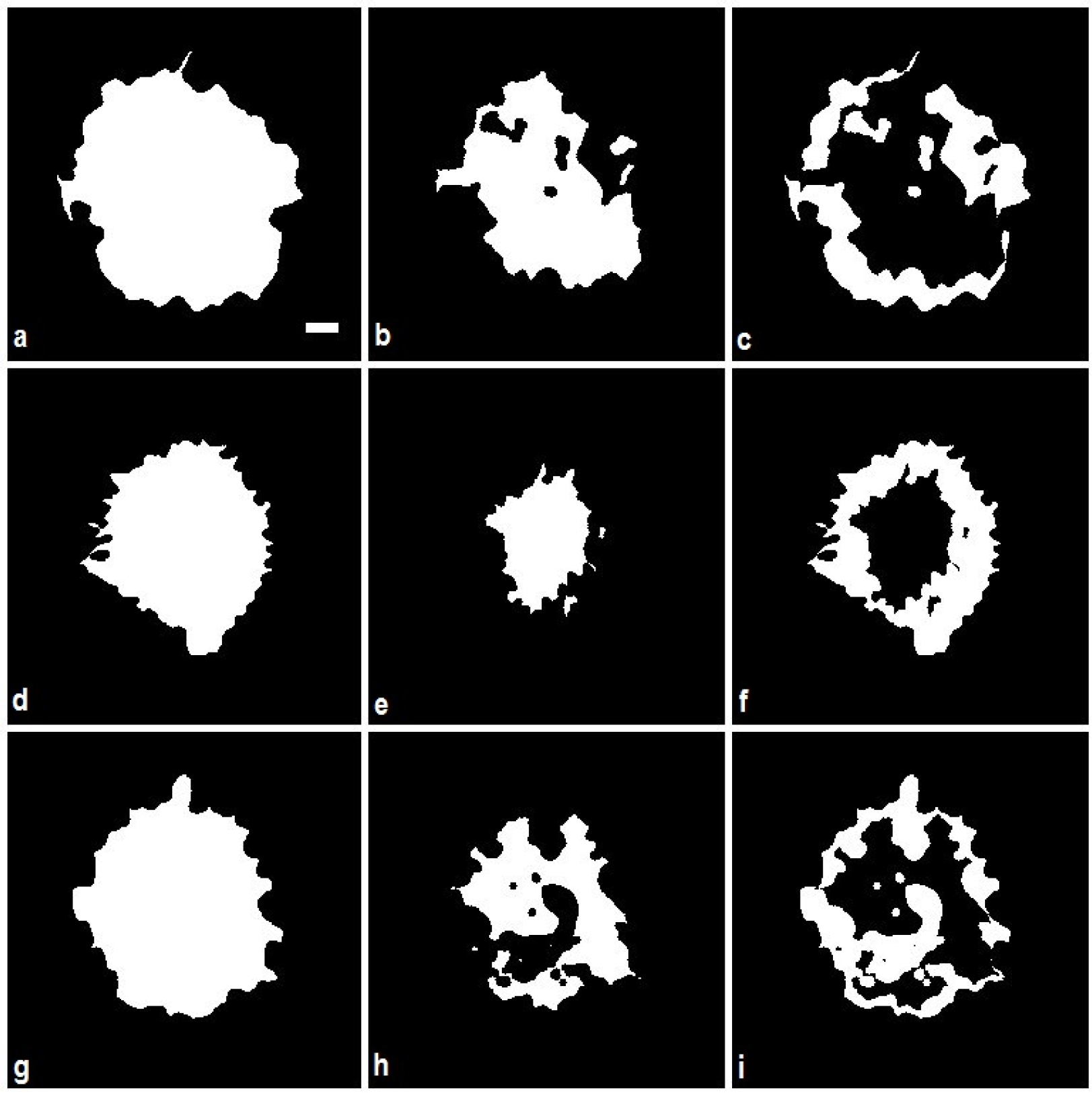
Image masks used to create normalized histograms of the number of pixels with particular P/S ratios in +IgE and −IgE regions shown in Figure 5d,h,l. Masks shown in the first row (panels a, b, and c) were used to create histogram shown in Figure 5d. Masks shown in the second row (panels d, e, and f) were used to create histogram shown in Figure 5h, and masks shown in the third row (panels g, h, and i) were used to create the histogram shown in Figure 5l. Masks shown in the first column represent cell masks obtained by thresholding the sum image of the P-polarized and S-polarized excitation. Masks shown in the second or middle column represent masks that contain IgE (+IgE) and were obtained by thresholding the IgE-488 image. Masks shown in the third column represent cell masks that lack IgE (−IgE) and were obtained by subtracting the corresponding +IgE masks from the cell masks in the first column. Scale bar shown in panel a) applies to all nine panels and represents 2 μm.

**Figure S2.**
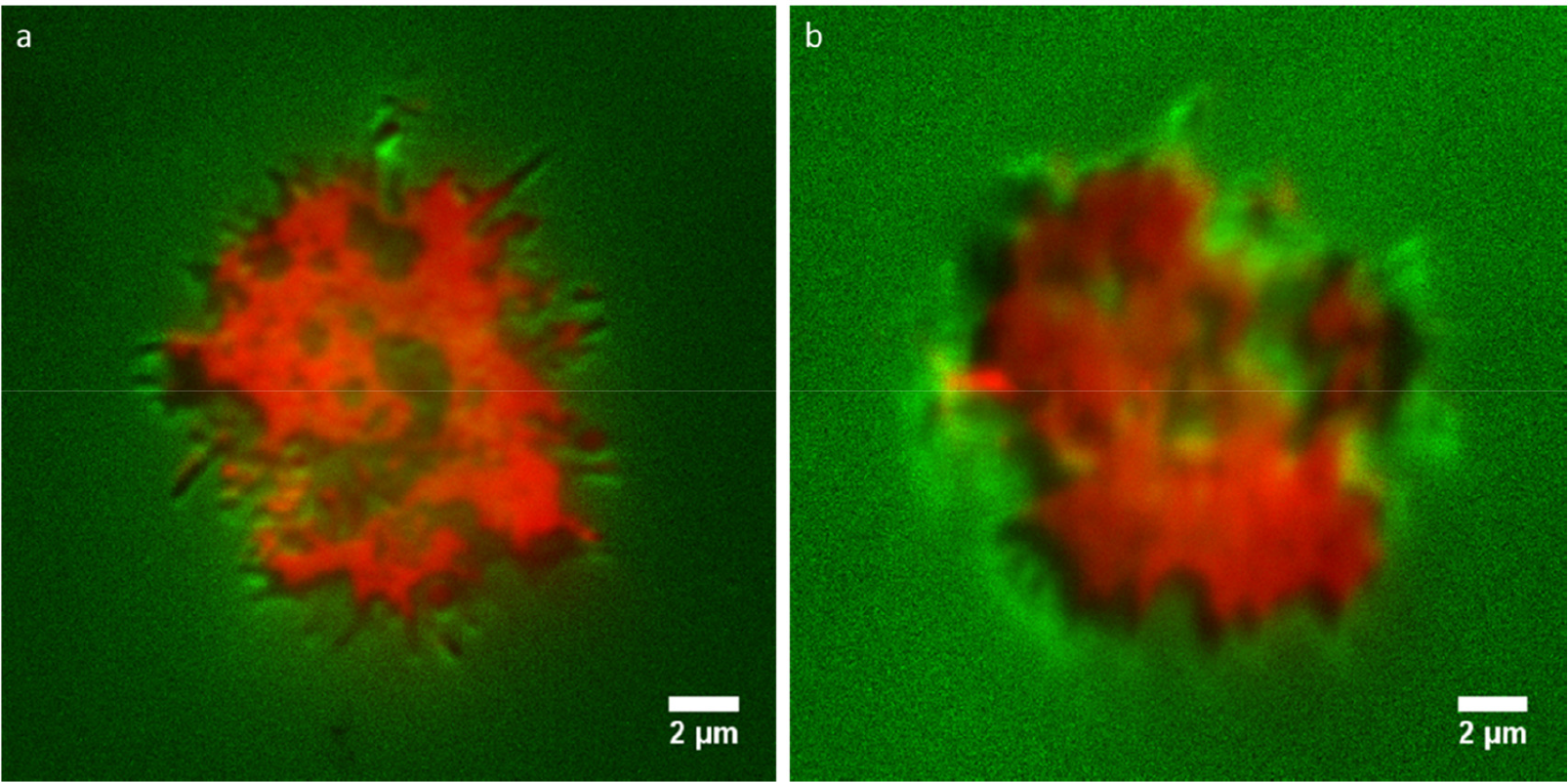
Image showing composite of IgE signal (red) and P-S ratio (green) from a) Figure 5i and 5k and b) Figure 5a and 5c of the main text.

## References

1. Monks, C.R.F.; Freiberg, B.A.; Kupfer, H.; Sciaky, N.; Kupfer, A. Three-dimensional segregation of supramolecular activation clusters in T cells. Nature 1998, 395, 82–86.

2. Kaizuka, Y.; Douglass, A.D.; Varma, R.; Dustin, M.L.; Vale, R.D. Mechanisms for segregating T cell receptor and adhesion molecules during immunological synapse formation in Jurkat T cells. Proc. Natl. Acad. Sci. 2007, 104, 20296–20301.

3. Carroll-Portillo, A.; Cannon, J.L.; Te Riet, J.; Holmes, A.; Kawakami, Y.; Kawakami, T.; Cambi, A.; Lidke, D.S. Mast cells and dendritic cells form synapses that facilitate antigen transfer for T cell activation. J. Cell Biol. 2015, 210, 851–864.

4. Carroll-Portillo, A.; Spendier, K.; Pfeiffer, J.; Griffiths, G.; Li, H.; Lidke, K.A.; Oliver, J.M.; Lidke, D.S.; Thomas, J.L.; Wilson, B.S.; et al. Formation of a mast cell synapse: Fc epsilon RI membrane dynamics upon binding mobile or immobilized ligands on surfaces. J. Immunol. 2010, 184, 1328–38.

5. Spendier, K.; Lidke, K.A.; Lidke, D.S.; Thomas, J.L. Single-particle tracking of immunoglobulin E receptors (FcεRI) in micron-sized clusters and receptor patches. FEBS Lett. 2012, 586, 416–21.

6. Spendier, K.; Carroll-Portillo, A.; Lidke, K.A.; Wilson, B.S.; Timlin, J.A.; Thomas, J.L. Distribution and dynamics of rat basophilic leukemia immunoglobulin E receptors (FcepsilonRI) on planar ligand-presenting surfaces. Biophys. J. 2010, 99, 388–97.

7. Song, J.; Hagen, G.M.; Roess, D.A.; Pecht, I.; Barisas, B.G. The mast cell function-associated antigen and its interactions with the type I FcE receptor. Biochemistry 2002, 41, 881–889.

8. Balakrishnan, K.; Hsu, F.J.; Cooper, A.D.; McConnell, H.M. Lipid hapten containing membrane targets can trigger specific immunoglobulin E-dependent degranulation of rat basophil leukemia cells. J. Biol. Chem. 1982, 257, 6427–33.

9. Thomas, J.L.; Feder, T.J.; Webb, W.W. Effects of protein concentration on IgE receptor mobility in rat basophilic leukemia cell plasma membranes. Biophys. J. 1992, 61, 1402–12.

10. Weis, R.M.; Balakrishnan, K.; Smith, B.A.; Mcconnell, H.M.; Smithy, B.A.; Mcconnell, H.M. Stimulation of fluorescence in a small contact region between rat basophil leukemia cells and planar lipid membrane targets by coherent evanescent radiation. J. Biol. Chem. 1982, 257, 6440–5.

11. Pfeiffer, J.R.; Seagrave, J.C.; Davis, B.H.; Deanin, G.G.; Oliver, J.M. Membrane and cytoskeletal changes associated with IgE-mediated serotonin release from rat basophilic leukemia cells. J. Cell Biol. 1985, 101, 2145–55.

12. Bassereau, P.; Jin, R.; Baumgart, T.; Deserno, M.; Dimova, R.; Frolov, V.A.; Bashkirov, P. V; Grubmüller, H.; Jahn, R.; Risselada, H.J.; et al. The 2018 biomembrane curvature and remodeling roadmap. J. Phys. D. Appl. Phys. 2018, 51, 343001.

13. Spendier, K. N-terminal amphipathic helix of Amphiphysin can change the spatial distribution of immunoglobulin E receptors (FcεRI) in the RBL-2H3 mast cell synapse. Results Immunol. 2016, 6.

14. Schmick, M.; Bastiaens, P.I.H. The interdependence of membrane shape and cellular signal processing. Cell 2014, 156, 1132–1138.

15. Axelrod, D. Carbocyanine dye orientation in red cell membrane studied by microscopic fluorescence polarization. Biophys. J. 1979, 26, 557–73.

16. Sund, S.E.; Swanson, J.A.; Axelrod, D. Cell membrane orientation visualized by polarized total internal reflection fluorescence. Biophys. J. 1999, 77, 2266–83.

17. Axelrod, D. Chapter 7: Total internal reflection fluorescence microscopy. Methods Cell Biol. 2008, 89, 169–221.

18. Passmore, D.R.; Rao, T.C.; Peleman, A.R.; Anantharam, A. Imaging plasma membrane deformations with pTIRFM. J. Vis. Exp. 2014.

19. Anantharam, A.; Axelrod, D.; Holz, R.W. Real-time imaging of plasma membrane deformations reveals pre-fusion membrane curvature changes and a role for dynamin in the regulation of fusion pore expansion. J. Neurochem. 2012, 122, 661–71.

20. Anantharam, A.; Axelrod, D.; Holz, R.W. Polarized TIRFM reveals changes in plasma membrane topology before and during granule fusion. Cell. Mol. Neurobiol. 2010, 30, 1343–9.

21. Anantharam, A.; Onoa, B.; Edwards, R.H.; Holz, R.W. Axelrod, D. Localized topological changes of the plasma membrane upon exocytosis visualized by polarized TIRFM. J. Cell Biol. 2010, 188, 415–28.

22. Johnson, D.S.; Toledo-Crow, R.; Mattheyses, A.L.; Simon, S.M. Polarization-controlled TIRFM with focal drift and spatial field intensity correction. Biophys. J. 2014, 106, 1008–19.

23. Oreopoulos, J.; Yip, C.M. Probing membrane order and topography in supported lipid bilayers by combined polarized total internal reflection fluorescence-atomic force microscopy. Biophys. J. 2009, 96, 1970–84.

24. Ramirez, D.M.C.; Jakubek, Z.J.; Lu, Z.; Ogilvie, W.W.; Johnston, L.J. Changes in order parameters associated with ceramide-mediated membrane reorganization measured using pTIRFM. Langmuir 2013,29, 15907–18.

25. Werner, J.H.; Montaño, G.A.; Garcia, A.L.; Zurek, N.A.; Akhadov, E.A.; Lopez, G.P.; Shreve, A.P. Formation and Dynamics of Supported Phospholipid Membranes on a Periodic Nanotextured Substrate. Langmuir 2009, 25, 2986–2993.

26. Edelstein, A.; Amodaj, N.; Hoover, K.; Vale, R.; Stuurman, N. Computer control of microscopes using μManager. Curr. Protoc. Mol. Biol. 2010, Chapter 14, 14.20.1–14.20.17.

27. Edelstein, A.D.; Tsuchida, M.A.; Amodaj, N.; Pinkard, H.; Vale, R.D.; Stuurman, N. Advanced methods of microscope control using μManager software. J. Biol. methods 1.

28. Friedman, L.J.; Chung, J.; Gelles, J. Viewing dynamic assembly of molecular complexes by multi-wavelength single-molecule fluorescence. Biophys. J. 2006, 91, 1023–31.

29. Ellefsen, K.L.; Dynes, J.L.; Parker, I. Spinning-spot shadowless TIRF microscopy. PLoS One 2015, 10, e0136055.

30. Hendriks, C.L.L.; van Vliet, L.J.; Rieger, B.; van Kempen, G.M.P.; van Ginkel, M. Dipimage: a scientific image processing toolbox for MATLAB; Quantitative Imaging Group, Faculty of Applied Sciences, Delft University of Technology, Delft, The Netherlands, 1999;

31. Pospíšil, J.; Lukeš, T.; Bendesky, J.; Fliegel, K.; Spendier, K.; Hagen, G.M. Imaging tissues and cells beyond the diffraction limit with structured illumination microscopy and Bayesian image reconstruction. Gigascience 2019, 8 giy126.

32. Gustafsson, M.G.L. Surpassing the lateral resolution limit by a factor of two using structured illumination microscopy. J. Microsc. 2000, 198, 82–87.

33. Ovesný, M.; Křížek, P.; Borkovec, J.; Švindrych, Z.; Hagen, G.M. ThunderSTORM: A comprehensive ImageJ plug-in for PALM and STORM data analysis and super-resolution imaging. Bioinformatics 2014, 30, 2389–2390.

34. Rust, M.J.; Bates, M.; Zhuang, X. Sub-diffraction-limit imaging by stochastic optical reconstruction microscopy (STORM). Nat. Methods 2006, 3, 793–795.

